# Formal Analysis of Network Motifs

**DOI:** 10.1101/347500

**Authors:** Hillel Kugler, Sara-Jane Dunn, Boyan Yordanov

## Abstract

A recurring set of small sub-networks have been identified as the building blocks of biological networks across diverse organisms. These network motifs have been associated with certain dynamical behaviors and define key modules that are important for understanding complex biological programs. Besides studying the properties of motifs in isolation, existing algorithms often evaluate the occurrence frequency of a specific motif in a given biological network compared to that in random networks of similar structure. However, it remains challenging to relate the structure of motifs to the observed and expected behavior of the larger network. Indeed, even the precise structure of these biological networks remains largely unknown. Previously, we developed a formal reasoning approach enabling the synthesis of biological networks capable of reproducing some experimentally observed behavior. Here, we extend this approach to allow reasoning about the requirement for specific network motifs as a way of explaining how these behaviors arise. We illustrate the approach by analyzing the motifs involved in sign-sensitive delay and pulse generation. We demonstrate the scalability and biological relevance of the approach by revealing the requirement for certain motifs in the network governing stem cell pluripotency.

## 1 Introduction

Network motifs [2, 19] are basic interaction patterns that recur throughout biological networks, where they are observed more frequently than in random networks with similar properties (e.g. a comparable number of components and interactions). The same small set of network motifs appears to serve as the building blocks of biological networks for diverse organisms [16, 31, 3]. Each network motif can operate as an elementary circuit with a well-defined function, which is integrated within a larger network and has a role in performing the required information processing [21]. Since the introduction of the concept of network motifs and the identification and experimental validation of initial instances [2], a wide range of additional motifs with new roles have been uncovered. Network motifs have been identified within transcriptional networks [26, 15], signaling networks [3], neuronal networks [24], and metabolic networks [23]. In addition to biological networks, recurring motifs have been identified in engineered systems, including electronic circuits and the world wide web [19].

The study of network motifs provides an attractive research direction towards understanding complex biological programs and uncovering modularity and reusable patterns of computation in the design of biological circuits. While many of the associated problems are challenging, especially when dealing with large biological networks, a wide range of computational methods have been developed [12, 28]. For example, novel motifs have been identified by comparing the occurrence frequency of a sub-network in a known biological network to that in random networks of similar structure, while various graph methods have been applied to algorithmically scan a network for specific motifs. Substantial research effort has been devoted to dealing with the algorithmic challenges of network motif identification and the related graph algorithms [17, 22, 1, 28]. This has led to the development of a number of computational tools, including mfinder [12], MAVisto [25], NeMoFinder [6], FANMOD [29], Grochow-Kellis [9], Kavosh [11], MODA [30], NetMODE [14], Acc-MOTIF [18] and QuateXelero [13] (see also [28] for a review and detailed comparisons). Verification techniques have also been applied to study network motifs. In [5], certain motifs and their dynamic properties were characterized using temporal logic, and parallel model checking was used to verify properties of networks with around ten components. In [10], approximate methods for analyzing gene regulatory networks were developed utilizing network motifs.

Following the identification of biologically-relevant motifs and the exploration of their dynamical properties in isolation, understanding how their presence or absence within a larger biological network defines that network’s behavior becomes a central problem. This problem is compounded by the fact that the precise structure of such biological networks often remains largely unknown, due to noisy and sometimes irreproducible experimental data. This makes it challenging to search for motifs within the network or to explore the connections between a network’s structure and its behavior.

Previously, we developed a formal reasoning approach enabling the synthesis and analysis of biological networks (e.g. incorporating gene regulation, signaling, etc.) that were only partially known [8, 32]. The method, summarized briefly in Sec. 2.1, introduced the concept of an Abstract Boolean Network as a formalism for describing discrete dynamical models of biological networks, where the precise interactions or update rules were unknown. These models could then be constrained with specifications of some required behaviors, thereby providing a characterization of the set of all networks capable of reproducing some experimental observations.

In this paper, we extend the approach from [8, 32] to enable automated reasoning about the requirement for specific network motifs as part of a biological network that is only partially known. This allows us to incorporate within the same framework constraints relating to the structure of the network, represented as logical formulas over the presence or absence of different motifs, together with constraints about the network’s dynamic behavior. Our reasoning approach then allows us to draw conclusions about certain motifs being essential or disallowed for reproducing the required behavior, thus helping explaining how the observed behaviors arise from various motifs.

We illustrate the approach by analyzing the motifs involved in sign-sensitive delay and pulse generation – two distinct behaviors that have been associated with certain network structures and observed biological properties [15, 16]. We consider a generic 3-layered network topology that serves as a prototype for a variety of biological programs and find that under the qualitative, Boolean modeling formalism of [8, 32], positive feedback is required to implement both behaviors.

To demonstrate the scalability and biological relevance of the approach, we apply the proposed method to identify the motifs requirements in the network governing stem cell pluripotency [8, 7]. This reveals that positive feedback, as well as a particular incoherent feed-forward motif, is also essential for maintaining pluripotency in the qualitative model.

We envision that the method proposed in this paper will provide a powerful tool for researchers interested in exploring the structural properties of biological networks and understanding how different motifs lead to various biological behaviors. In the future, this tool could support theoretical studies, where the connections between network structure and function are explored, as well as experimental studies, for example allowing researches to focus on the core, essential modules of biological networks.

## 2 Methods

In the following, we introduce some notation and summarize the approach from [32], which is implemented in the computational tool RE:IN (Sec. 2.1) and serves as a foundation for the extensions we propose in this paper (Sec. 2.2 and Sec. 2.3).

### 2.1 Abstract Boolean Network Analysis

Following the notation from [32], an *Abstract Boolean Network (ABN)* is a tuple 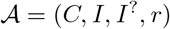, where

– *C* is the finite set of components,
– *I* is the set of definite (positive and negative) interactions between the components from *C*,
– *I*^?^ is the set of possible (positive and negative) interactions, and
– *r* assigns a subset of regulation conditions (possible update functions) to each component from *C*.

ABNs are discrete, dynamic models suitable for studying biological systems, when often the existence of interactions between components are hypothesized, but not definitively known [8, 32]. An example of an ABN is illustrated in Fig. 1a.

**Fig. 1.**
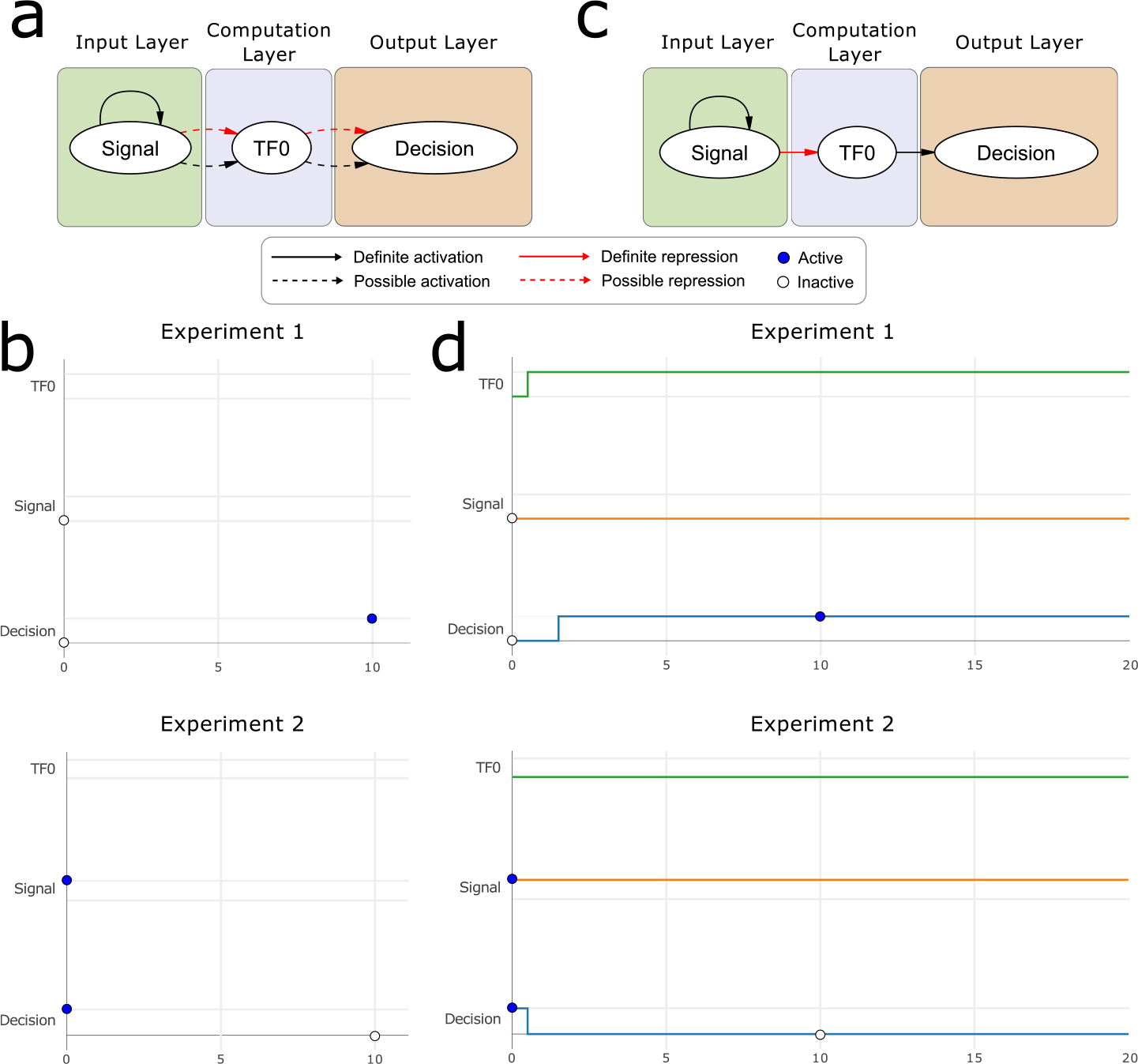
Abstract Boolean Networks constrained against experimental observations. **a)** The generic network architecture we consider is comprised of three layers (Input/Computation/Output). The input and output layers each include a single component, while the number of computation components *n* can be varied (in this case *n* = 1). This simple ABN includes 4 optional interactions and 1 definite interaction. Formally, the ABN is defined as 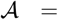 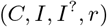, where *C* = {`Signal`, `TFO`, `Decision`}, *I* = {(`Signal`, `Signal`, +)}, *I*^?^ = {(`Signal`, `TFO`, +), (`Signal`, `TFO`, −), (`TFO`, `Decision`, +), (`TFO`, `Decision`, −)}, and *r* allows all regulation conditions for each component. **b)** Experimental constraints encode expected states along different network trajectories. Here, two experimental constraints are illustrated, which specify the initial state of the signal and decision components, and their state at step 10. **c)** A single, concrete network that is consistent with the constraints in (b) is generated using our SMT-based approach. **d)** Experiment trajectories from the concrete model in (c) illustrate how the experimental constraints are satisfied by this network.

Each component *c* ∈ *C* describes a different chemical signal, protein, gene, etc. that can exist in one of two states: active or inactive. The dynamics of the system are defined by the regulation condition assigned to each component, which serves as an ‘update function’ that specifies the state of the component at step *k* + 1, given the state of all of its regulators (other components *c*′ ∈ *C* with interactions to *c*) at step *k*. Here we consider synchronous updates, such that deterministic trajectories emerge from each initial state, though RE:IN also allows the exploration of asynchronous trajectories.

ABNs are abstract models because of the uncertainty in the precise network topology and regulation rules for each component. An ABN is transformed into a concrete Boolean Network by instantiating a subset of the possible interactions, discarding all other optional interactions, and assigning a specific regulation condition for each gene. By virtue of the unique combination of interactions and regulation conditions, different concrete models derived from the same ABN can have different dynamical behaviors.

The concept of a *Constrained Abstract Boolean Network (cABN)* was introduced in [32] as a formalism for describing a set of Boolean Network models that are consistent with some experimentally observed biological behaviors. A cABN is defined in terms of an ABN, together with a set of constraints over the states of the components from *C*. These constraints encode experimental observations, where separate executions of the system correspond to different biological ‘experiments’. For example, the observations encoded in Fig. 1b specify a biological program where cells make a particular decision only in the absence of some signal. Experiment 1 requires that there exists a trajectory, where both the ‘Signal’ and ‘Decision’ components are initially inactive and ‘Decision’ becomes active at step 10. Similarly, Experiment 2 requires that there exists a trajectory, where both the ‘Signal’ and ‘Decision’ components are initially active and ‘Decision’ becomes inactive at step 10. These constraints limit the feasible assignments of regulation conditions and possible interactions, such that all concrete networks from the cABN are guaranteed to reproduce all experimental observations.

cABN analysis was solved in [32] by encoding it as a Satisfiability Modulo Theories (SMT) problem. This enables the enumeration of individual concrete models that are consistent with the experimental observations. For example, the concrete model from Fig. 1c is generated from the ABN in Fig. 1a and is consistent with the constraints from Fig. 1b. This is demonstrated using the trajectories visualized in Fig. 1d. The SMT-based approach from [32] also allows reasoning about hypotheses describing unknown biological behaviors to make novel predictions from all consistent models collectively, without the need to enumerate individual concrete networks.

In addition to the above, the analysis can reveal both required and disallowed interactions of the cABN. An interaction *i* ∈ *I*^?^ is required if the experimentally observed behavior cannot be reproduced without it (i.e. all concrete models of the cABN include the interaction *i*). Similarly, an interaction is disallowed if including it in a concrete model means that the observed behavior can no longer be reproduced. The analysis of required and disallowed interactions yields insight into how network structures lead to certain dynamic behaviors, and is the starting point for the extensions we propose here.

In the following, we use set notation to denote the existence or non-existence of interactions in an ABN or a cABN. For example, (*c*, *c*′, +) ∈ *I* denotes that a definite, positive interaction exists in 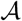, (*c*, *c*′, −) ∈ *I*^?^ denotes that an optional, negative interaction exists, and (*c*, *c*′, ∗) ∉ *I* denotes that no definite interactions (i.e. the wild card ∗ stands for either + or −) exist between *c* and *c*′.

### 2.2 Motif Assignment

#### Definition 1 (Motif).

*A motif is a tuple* 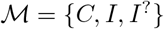, *where C is the finite set of components, I is the set of definite and I*^?^ *the set of possible interactions (similarly to the definition of ABNs).*

Examples of several different motifs are illustrated in Fig. 3. In contrast to ABNs, motifs (Def. 1) are static networks, without regulation conditions (update functions) to make them dynamical systems. However, because interactions from *I*^?^ are uncertain, motifs are abstract - a motif defined as in Def. 1 with a nonempty *I*^?^ describes a set of 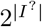 concrete, static networks.

#### Definition 2 (Motif Assignment).

*Given an ABN* 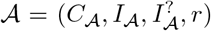 *and a motif* 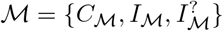 *a motif assignment is a map* 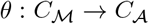.

Note that Def. 2 can also be applied to cABNs instead of ABNs by omitting the set of experimental observations from the cABN.

Given an ABN 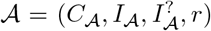 and a motif 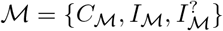, let 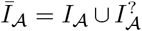 and 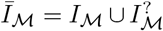 denote the set of all interactions (definite and optional) in the ABN and the motif. Given a motif assignment 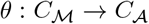, let 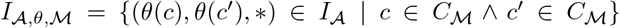 denote the set of definite interactions from the ABN between components that the motif maps to. Similarly, let 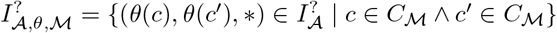 denote the set of optional interactions from the ABN between components that 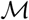 maps to^5^.

#### Definition 3 (Valid Motif Assignment).

*A given motif assignment θ between ABN* 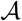 *and motif* 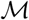 *is valid if and only if*

1. 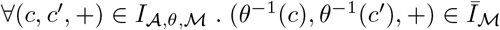,
2. 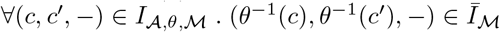,
3. 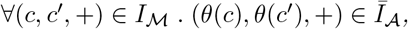, and
4. 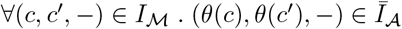

The conditions from Def. 3.1-3.4 ensure that the motif components are assigned to ABN components in such a way that the interactions match. In other words, each definite (positive or negative) interaction in the ABN (between components that the motif maps to) matches an interaction (definite or optional) in the motif (Def. 3.1-3.2) and each definite interaction in the motif matches an interaction in the ABN (Def. 3.3-3.4).

Given an optional interaction 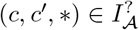, let 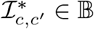 denote the Boolean choice variable representing whether the interaction is included in a concrete model (see [32] for details of the SMT encoding of ABNs and cABNs). Asserting that 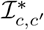 is true can be interpreted as modifying the ABN such that 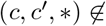 and 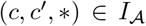 (i.e. ensuring that the interaction is definitely present). Similarly, asserting that 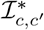 is false can be interpreted as modifying the ABN such that 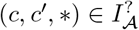 but 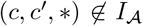 (i.e. ensuring that the interaction is definitely absent).

The notion of a valid motif assignment (Def. 3) is sufficient to guarantee that the components of the motif are mapped to components of the ABN in such a way that all definite interactions are matched. However, it is possible that optional interactions of the ABN map to definite interactions of the motif or do not match any motif interactions. Therefore, while the interactions of the ABN match that of the motif, it is not possible to guarantee that every concrete network represented by the ABN matches the motif. The additional constraints defined in the following ensure that this is indeed the case.

#### Definition 4 (Motif Assignment Constraints).

*Given a motif assignment θ between ABN* 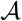 *and motif* 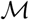, *the motif assignment constraints are*

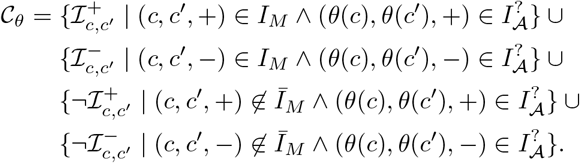

The additional constraints from (Def. 4) assert that an optional interaction in the ABN that matches a definite interaction of the motif is always included. Similarly, an optional ABN interaction that does not match any motif interaction is never included. These additional constraints guarantee that the interactions of all concrete networks of the ABN match those of the motif, under the given motif assignment.

### 2.3 Motif Constraints

The motif assignment constraints (Def. 4) ensure that a given motif 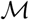 is implemented in all concrete networks of an ABN 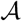 between the specific components defined by the motif assignment *θ*. In general, however, we are interested in guaranteeing that motif 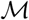 is implemented in the ABN 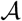 by any of its components rather than the the specific set of components specified by *θ*.

#### Definition 5 (Motif Constraints).

*Given an ABN* 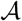 *and a motif* 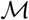, *the motif constraints* 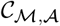 *are defined in terms of the motif assignment constraints (Def. 4) as* 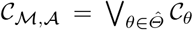, *where* 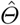 *is the set of valid motif assignments between* 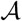 *and* 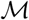 *(Def. 2)*.

The motif constraints from Def. 5 guarantee that motif 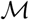 is implemented by every concrete network of ABN 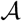, even though the precise components used to implement 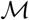 might differ.

Given an ABN 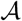 and a set of motifs, logical formulas (e.g. 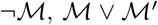, 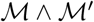, etc) could be constructed and interpreted by replacing each motif 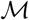 with its corresponding motif constraints 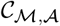.

### 2.4 Implementation

As part of the RE:IN framework [8, 32], a high-level, domain specific language was proposed for describing cABNs by defining the sets of components and interactions, as well as associated experimental observations. We implement the methods described in Sec. 2.2 and Sec. 2.3 as an extension of RE:IN that enables the reasoning about cABNs with additional structural constraints about the presence or absence of various motifs.

Currently, the generation of motif constraints (Def. 5) is implemented as a pre-processing step using a straightforward, exhaustive algorithm, where all motif assignments are first generated and then filtered to preserve only the valid ones using the conditions from Def. 3. Various cABN analysis problems are then encoded and solved using an SMT solver as shown previously [8, 32], while the additional motif constraints are also incorporated.

Two notable modifications are introduced to the method described in Sec. 2.2 and Sec. 2.3 for improved usability. First, when the name of a motif component matches the name of a cABN component, no other assignments are considered for that component. This enables the specification of partially known motifs, where the mapping of some of the motif components to the cABN components is given. Second, a dummy ‘Context’ component is always included within the set of motif components. Given a motif assignment, the ‘Context’ component matches any cABN component that is not already mapped to by the motif. This provides additional control in specifying how a motif could be implemented as part of the cABN’s network. For example, including optional positive and negative interactions from ‘Context’ to every motif component and vice versa does not impose additional constraints on the motif’s implementation. Without any additional ‘Context’ interactions, on the other hand, the motif can only be fully isolated and disconnected from all other components of the cABN.

### 2.5 Reasoning about Motifs

Combining the previously-developed SMT-based reasoning strategies [8, 32] with an encoding of the motif constraints from Sec. 2.3 enables the automated reasoning about structural (motif) properties of a network, together with the requirements about reproducing certain dynamical behaviors. Among the different analysis questions this method could support, in this work we focus specifically on identifying required (essential) and disallowed motifs. Similarly to the identification of required and disallowed interactions (Sec. 2.1), a motif 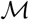 is required if the experimentally observed behavior could not be reproduced without it, while it is disallowed if enforcing that the motif is present in the network guarantees that the observed behavior can no longer be reproduced. These hypotheses can be tested as follows. Applying the SMT analysis of an ABN, if no concrete models are identified with the constraint 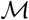 (the motif is present in the cABN) then the motif is disallowed. If, on the other hand, no concrete models are identified with the constraint 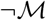 (the motif is not present in the network), then the motif is essential.

## 3 Results

To illustrate the analysis method proposed in Sec. 2, we apply it to study the importance of certain motifs for biological networks. First, we consider a generic network architecture (Fig. 2a) composed of an input, computation, and output layer, which serves as a prototype for many biological programs (Sec. 3.1). We study the models consistent with this network topology that could give rise to two distinct dynamical behaviors (Fig. 2b) and identify the motifs from a given set (Fig. 3) that are required or disallowed for producing this behavior as described in Sec. 2.5. Then, in Sec. 3.2 we apply our motif analysis method to the recently identified biological program governing stem cell decision making [7], demonstrating the scalability and biological relevance of the approach.

### 3.1 Biological Program Prototype

We construct a simple abstract network topology in order to explore how various motifs give rise to different dynamical behaviors. The network has a single input component that represents some biochemical signal and a single output (readout) component. The output component might represent a biochemical signal that affects a downstream process or, as in this example, could represent a particular cellular decision (e.g. to differentiate, divide, etc). Information processing is performed in the ‘computation’ layer, which includes a number of components. While all interactions in the network are unknown, we assume that information flows from the input layer, through the computation layer, and into the output layer. As a result, we consider a network with a densely connected computation layer (possible positive and negative interactions between each pair of computation components). The input (signal) component might affect any of the computation components, so possible positive and negative interactions from the input (signal) to all computation components are included. Similarly, possible positive and negative interactions from each computation component to the output (decision) component are included. A definite self-activation is included for the signal to guarantee that once set at the beginning of computation, its value does not change, but no other self-regulation interactions are allowed. The resulting network architecture with *n* = 3 computation components is visualized in Fig. 2a. In all subsequent analysis, we impose the additional constraint that a positive and a negative interaction between the same components are never included together in concrete models.

**Fig. 2.**
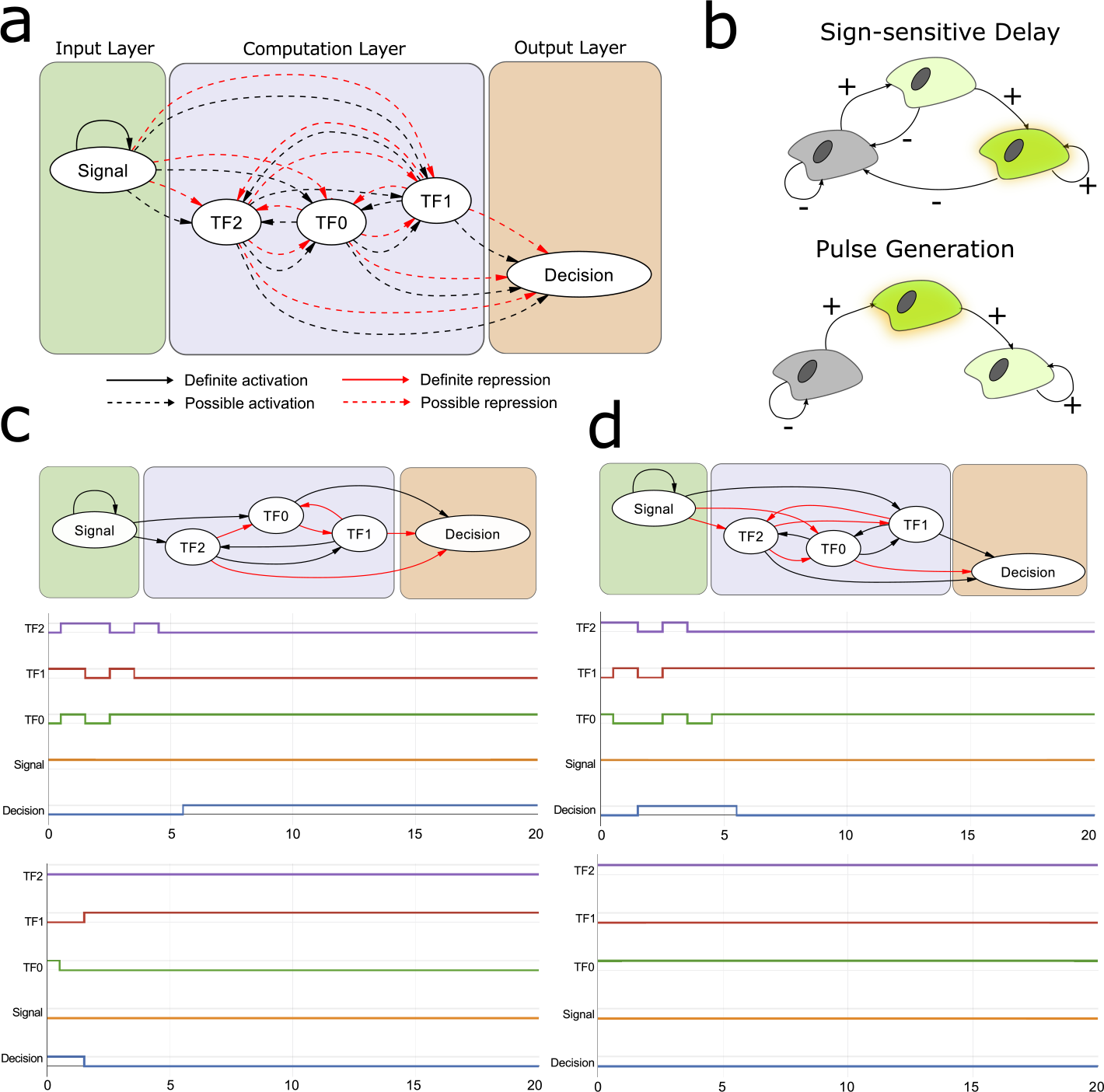
A generic network architecture generates biological programs implementing either sign-sensitive delay or pulse generation. (**a**) The ABN we consider is comprised of three computation components, with optional positive and negative interactions between them. (**b**) The sign-sensitive delay and pulse generation behaviors are represented graphically as transitions between different cellular states. Signs on the edges indicate the presence (+) or absence (−) of input signal, while the output is active only in the cell type shown in green (right-most cell). (**c**) An example of a biological program implementing sign-sensitive delay has the characteristic delay during activation (top trajectories) but responds faster during deactivation (bottom trajectories). (**d**) An example of a biological program implementing pulse generation produces a single pulse of the output when the input signal is present (top trajectories), while the output remains inactive when the signal is not present (bottom trajectories).

**Fig. 3.**
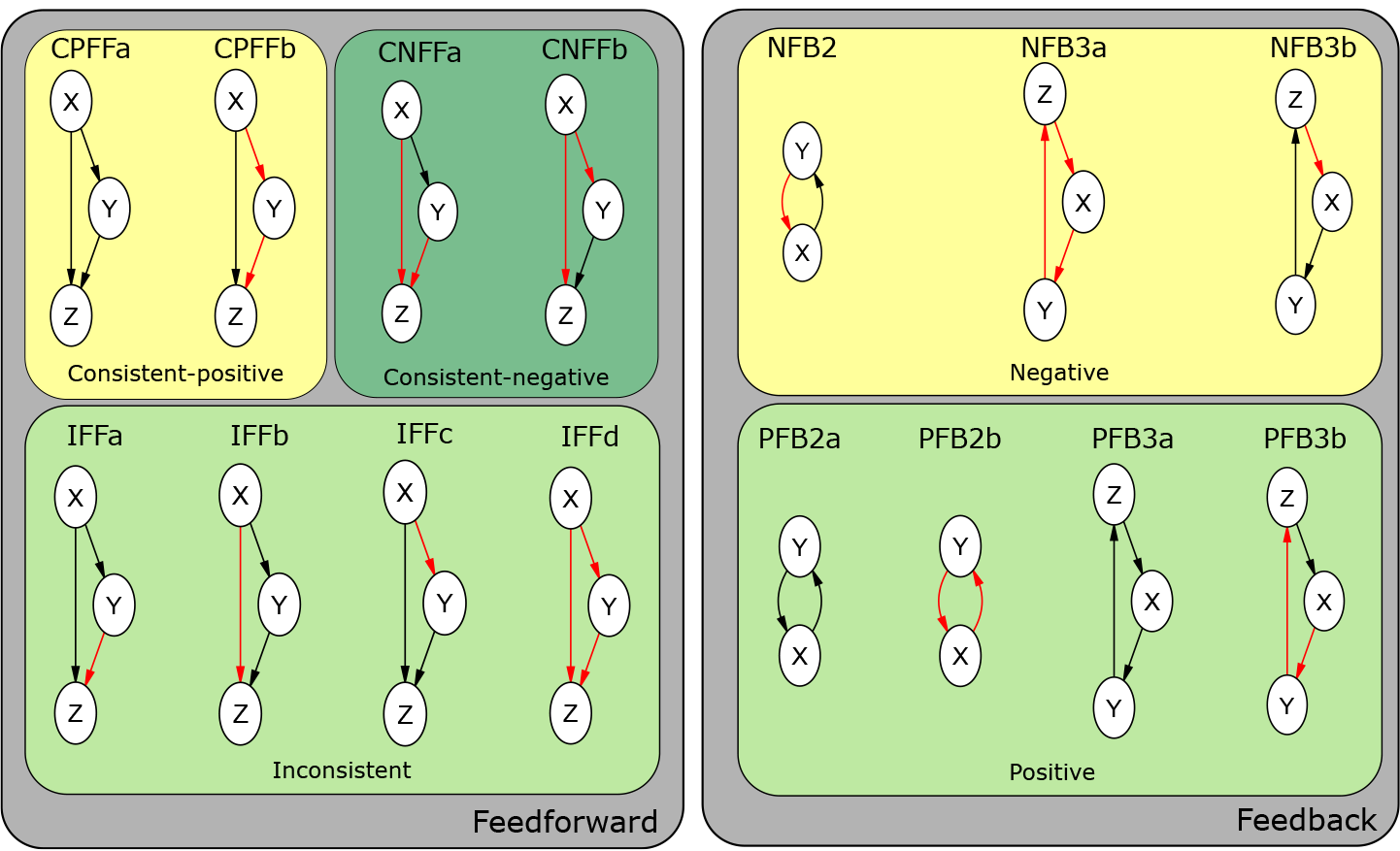
The set of 15 network motifs we define and use to analyze biological programs. These are sub-divided into a set of feedforward and feedback motifs. An optional positive and negative interaction from the ‘Context’ component to every motif component, and from every motif component to ‘Context’, is also included but not shown in the figure. This ensures that the motif of interest can be identified in the ABN being explored regardless of how the motif components are connected within the network external to the motif (see Sec 2.4).

#### Sign-sensitive delay

The first dynamical behavior we consider requires that a system produce an output in response to some input (e.g. the ‘Decision’ component becomes active if and only if ‘Signal’ is present in Fig. 2a). However, while the effect on the output is immediate when the signal is withdrawn, there is a delay on the output’s activation when the signal is supplied [^6^ Depending on the exact implementation, the delay can be observed when the signal switches from active to inactive instead, but this variation of a sign-sensitive delay is not considered here.]. Due to the asymmetric response to changes in the input signal, the behavior is called sign-sensitive delay [15, 16] and was shown to have a potential role for making decisions based on noisy inputs by filtering out fluctuations in input stimuli [15, 16].

We encode the requirement for a sign-sensitive delay as depicted in Fig. 2b. When no signal is present, the system can stabilize in a state where the output is inactive (shown in gray). When the signal is supplied, a transition to an intermediate state occurs (light green), although the output is still not activated. Withdrawing the signal at this point resets the system to the initial state, while continuous application of the signal leads to a transition to a state where the output is active (green). This active state is stable as long as the signal is present, but withdrawal of the signal leads quickly back to the initial, inactive state. The number of steps before the output is activated upon supplying the signal is the delay.

We find that for delays greater than a single step, a network with at least *n* = 3 computation components is required, and with *n* = 3, delays of up to 4 steps can be produced. An example of a concrete network implementing this behavior is shown in Fig. 2c, but many such networks consistent with the cABN from Fig. 2a exist, involving a variety of network motifs. To investigate further the network structures capable of producing sign-sensitive delays, we defined a number of feed-forward and feed-back motifs shown in Fig. 3.

We found that, while several motifs (**IFFa**, **IFFc-d**, **PFB3a-b**, **NFB3a-b**) were disallowed (possible due to the limited number of computation nodes), no motifs were required for producing a sign-sensitive delay of 4 steps (see Table 2). We then considered pairs of motifs (e.g. by testing 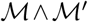 and 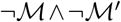) and found that, while many pairs were jointly disallowed, only the pair **PFB2a** and **PFB2b** was jointly required (see Table 2). This indicates that a positive feedback motif between two components (either **PFB2a** or **PFB2b**) is essential for implementing a 4-step sign-sensitive delay in a network with *n* = 3 computation components, but one of these motifs can be substituted for the other.

#### Pulse generation

The second dynamical behavior we consider requires that a system produce a transient output pulse in response to some input. We encode this behavior as depicted in Fig. 2b. While no signal is present, the system remains stably in a state where the output is inactive (shown in gray). When the signal is supplied, a transition to an intermediate state occurs (green) where the output is activated. Further application of the signal causes a transition to a state where the output is no longer active (light green). Currently, no ‘resetting’ (i.e. withdrawal of the signal causing a transition to the initial state) is considered. The number of steps during which the output is active upon supplying the signal is the pulse width.

We find that for pulse widths greater than a single step, a network with at least *n* = 3 computation components is required, and with *n* = 3, pulse widths up to 4 steps can be produced. An example of a concrete network implementing this behavior is shown in Fig. 2d.

As in the previous case study, we were interested in exploring how the motifs from Fig. 3 affect the capacity for pulse generation. We found that, several motifs (**CPFFa**, **IFFc-d**, **PFB3a-b**, **NFB3a-b**) were disallowed and **PFB2b** was required. This indicates that a positive feedback motif between two components, implemented specifically through two negative interactions, is essential for implementing a generator for pulses of width 4 in a network with *n* = 3 computation components.

### 3.2 Stem Cell Pluripotency Program

The sign-sensitive delay and pulse generation examples demonstrate the utility of our approach in revealing how different network motifs give rise to different dynamic behaviors within a relatively simple network. Here we apply our analysis to a realistic biological network. Previously, we used RE:IN to study the biological program governing maintenance of naïve pluripotency: the property uniquely exhibited by embryonic stem cells (ESCs) to generate all adult cell types, as well as the germline. This capacity is lost as cells begin to differentiate into the different adult lineages throughout embryogenesis. While the critical transcription factors (TFs) that regulate pluripotency had been identified, and the culture environments required to sustain the cells progressively refined [20], it was not previously understood how environmental signals were processed by the core TFs to govern the pluripotent state. We investigated the biological program governing pluripotency by defining an ABN based on gene expression profiles of mouse ESCs, which was subsequently constrained against a rich set of behaviors derived from experiments in which ESCs are subject to different culture conditions and molecular perturbations [8]. This network has since been further refined with additional data [7] (Fig. 4a). It is of interest to learn from this complex starting set of networks whether there are specific motifs that are required to generate, and thereby explain, the experimental observations of pluripotency.

Our analysis revealed that three motifs are required in all concrete models, **IFFa**, **PFB2a**, **PFB3a**, and four are disallowed, **CNFFa**, **IFFc**, **NFB2**, **NFB3a**. The set of disallowed motifs is a trivial result, as none of these are present in the ABN, regardless of how the optional interactions are instantiated. In contrast, there are a number of possible configurations of components in the ABN that are equivalent to the three motifs identified as required. We subsequently tested the requirement for specific component assignments of these motifs, and found that there are four cases present in all concrete models that are consistent with the experimental constraints. The four required motifs are shown in Fig. 4b, revealing that these models all require both feed-forward and feed-back motifs between specific network components.

This example demonstrates how our approach scales to complex networks that explain critical aspects of cellular decision-making, and can reveal essential elements of these biological programs required to explain observed behavior. Further analysis could reveal whether the motifs we have identified serve to determine the robustness of the pluripotency network, which is known to vary between different culture conditions [8]. Furthermore, it would be of interest to study whether these required motifs are repeated throughout networks governing similar biological behaviors, and in particular, whether they arise in pluripotency networks from other mammals, revealing elements of pluripotency that are conserved between species.

## 4 Discussion

The SMT-based formal reasoning approach from [8, 32] allows us to encode dynamic Boolean models of biological networks, where the precise set of interactions and regulation conditions for each component are unknown. These models can be constrained against specifications of experimental observations in order to identify concrete models capable of reproducing the required behavior, or to test novel hypotheses without selecting a particular concrete model from the set of models consistent with the experiments.

**Fig. 4.**
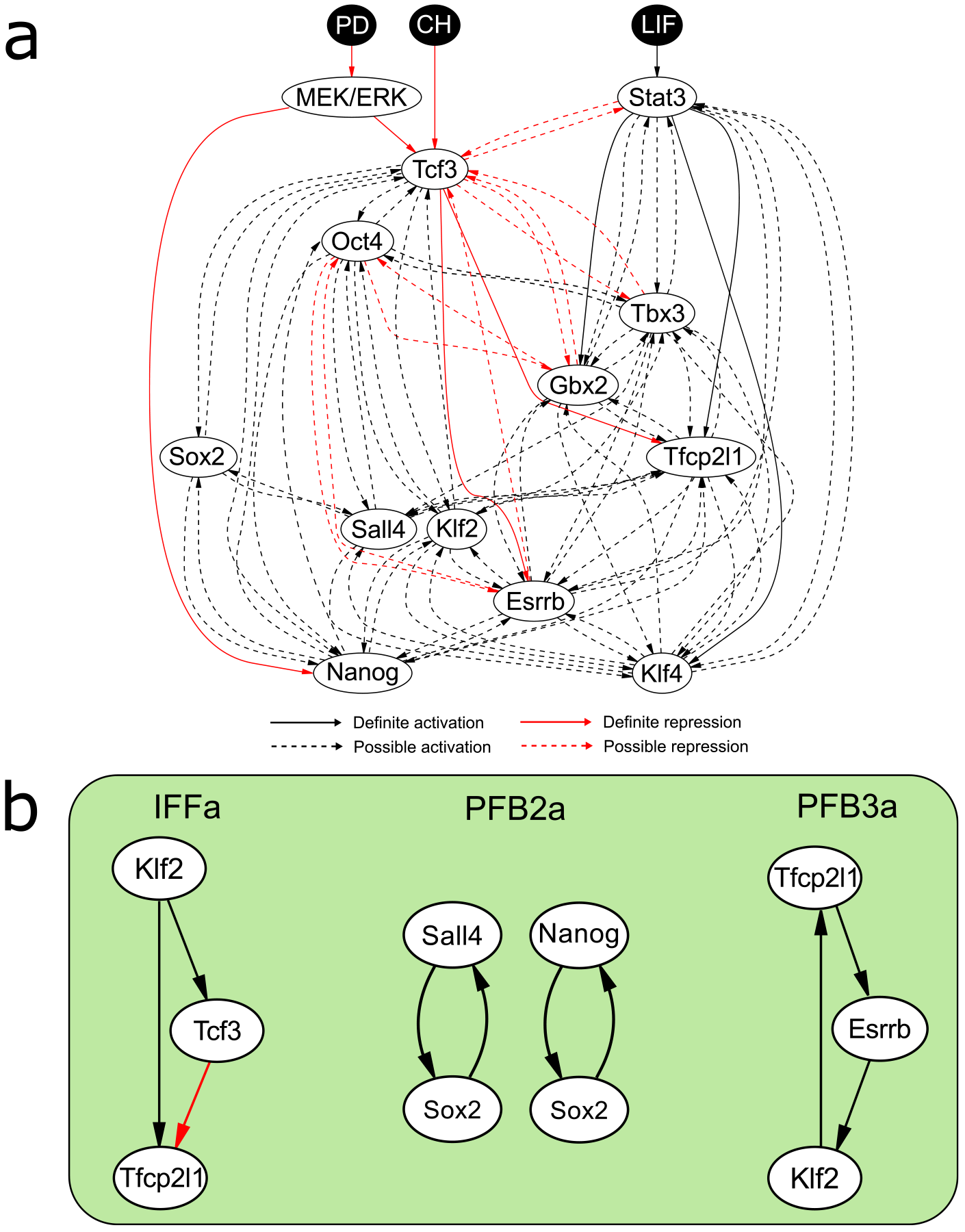
Network motifs required for stem cell pluripotency. (**a**) The ABN defined for three input signals provided as culture conditions to mouse ESCs - LIF, CH and PD - and 13 downstream components that have been functionally validated as critical pluripotency factors [7]. (**b**) Four instantiations of the three motifs required to satisfy the experimental constraints.

Certain structural constraints can also be handled directly by the previous method. For example, assigning an interaction as definite guarantees that only concrete models that incorporate that interaction are considered. Similarly, removing an interaction as either optional or definite guarantees that none of the considered models incorporate this interaction. However, expressing more complex structural properties, such as those required for reasoning about motifs, is challenging using the approach from [8, 32]. The extensions we propose in this work allow for complex properties describing the presence or absence of arbitrary motifs to be specified and tested, even when the network being studied is partially unknown. This leads to a natural framework for incorporating rich structural constraints and jointly reasoning about motifs and dynamical properties.

While the problem of identifying motifs does not scale favorably to large networks [4, 27], in [8] it was demonstrated that relatively small networks of core components can explain a rich set of biologically-relevant behaviors and cellular decisions. Therefore, even the straightforward, exhaustive algorithm implemented currently to generate motif constraints (Sec. 2.4) proves suitable for examining networks of biological significance. Indeed, the computation times for generating motif constraints are negligible compared to the task of verifying whether any concrete models exist that satisfy all constraints (see results in Appendix A). All analysis reported in this paper was accomplished on the order of seconds to minutes for simple networks (Sec. 3.1) and tens of minutes for the biological network governing stem cell pluripotency on a 3.6GHz Intel Xeon (E5-1620) computer with 32GB RAM (Sec. 3.2). Even so, many of the algorithmic advances towards more efficient identification of motifs in large networks [17, 12, 28] could be adapted as part of the pre-processing step of motif constraint generation.

An obvious limitation of the proposed method is that the cABN models being analyzed are qualitative (Boolean), discrete, and also deterministic, due to the synchronous update semantics we assume (although asynchronous updates are also supported by the method from [8, 32]). Therefore, dynamical behaviors associated with motifs or networks that require more detailed modeling assumptions cannot be handled directly. Thus, certain properties, for example relating to noise propagation and attenuation, or precise timing of signals, are not currently supported by our motif analysis.

Still, a number of interesting biological questions can be framed in terms of the analysis of structural, motif-based properties of partially known networks with respect to the dynamical behaviors they produce. In this work, we focused specifically on identifying motifs that are essential or disallowed for producing certain behaviors. The proposed approach can also be used for more in-depth studies, for example in order to identify whether an essential motif must always involve specific components in the network, as illustrated in the stem cell case study from Sec. 3.2. This is achieved by testing whether concrete models without a given motif involving a particular network component exist. Further detailed studies could also explore not just the presence of absence of motifs, but also how these motifs must be connected to the rest of the network (e.g. for a particular flow of information between components) in order to achieve the required behavior by exploring different combinations of optional and definite interactions between motif components and the ‘Context’.

Another interesting question is what additional insights the motif analysis proposed in this work can provide beyond the identification of required and disallowed interactions, already possible with the methods from [8, 32]. Intuitively, if a given motif must be implemented by specific components of a network, then all the motif’s interactions would be identified as required, which is the case for some of the motifs from the stem cell case study (Sec. 3.2). However, identifying motifs that are essential but appear in different places in different concrete networks that are consistent with experimental observations, reveals deeper connections between structure and behavior. The degree to which a network is constrained, either by reducing the number of optional interactions or by specifying additional behaviors that must be reproduced, limits the possibility of implementing motifs between different components and could lead to more predictions about essential or disallowed motifs. In contrast, larger, less-constrained biological networks can achieve the same behavior in different ways and certain motifs would no longer be required.

The case studies presented here illustrate how the proposed method can be used for theoretical studies of the properties and requirements for different motifs (Sec. 3.1) and demonstrate that the approach scales up to and provides useful insights about realistic biological networks (Sec. 3.2). Extending these studies and experimentally validating the motif requirements in these networks is a direction for future research.

## 5 Summary

To deal with the challenge of studying the connections between the structure and function of biological networks, even when these networks are only partially understood, we extended the SMT-based reasoning approach RE:IN from [8, 32]. The proposed methods involve the algorithmic generation of motif constraints, encoding the requirement that a given motif is present in some cABN. Rich structural requirements can then be incorporated in addition to the functional properties encoded as part of the cABN through logical formulas over such motif constraints. We illustrated the method by predicting that certain motifs are essential or disallowed for producing sign-sensitive delay and pulse generation in a network representing a prototype of biological programs, and in the biological network governing stem cell pluripotency [7]. The proposed methods enable the study of network motifs in the context of partially unknown, abstract networks and can support future theoretical and experimental studies, where reasoning about network structure and function in the same framework is essential.

## A Appendix: Detailed Analysis Results

**Table 1.**
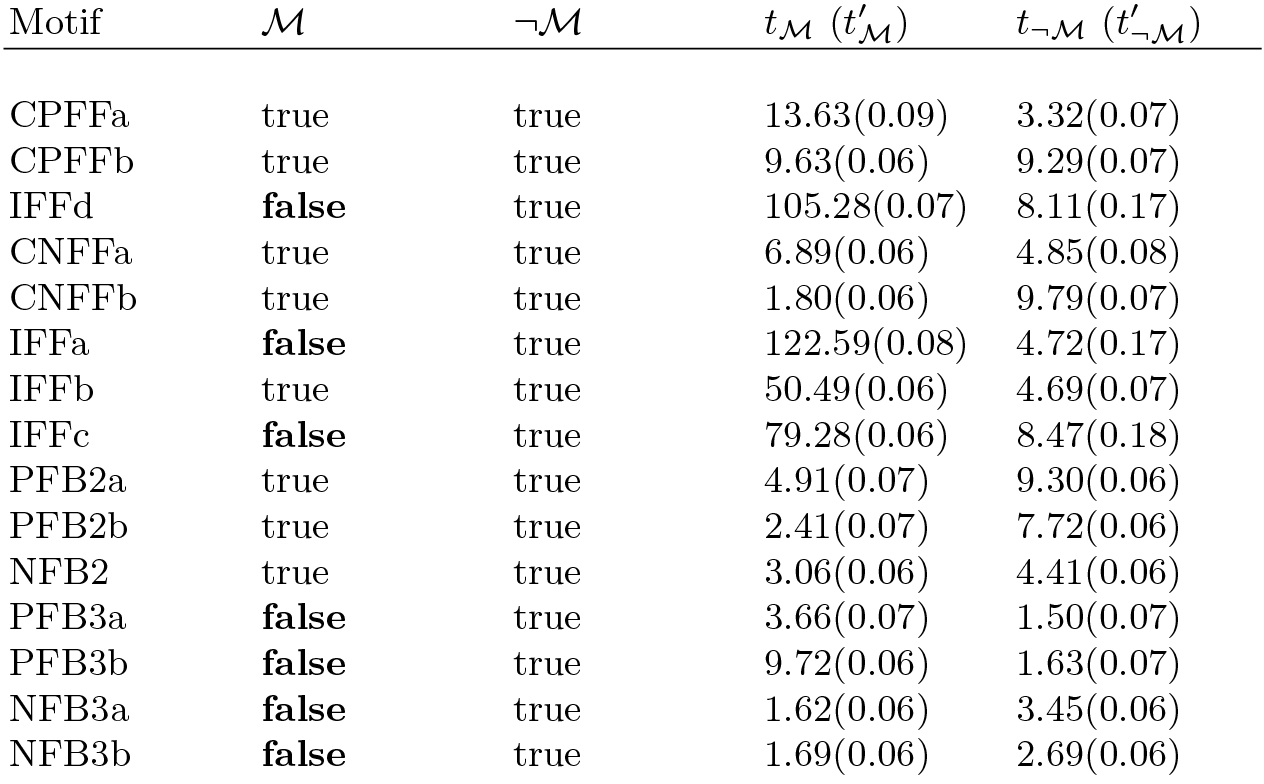
Single-motif analysis results for the 4-step sign-sensitive delay property of the *n* = 3 generic network. Column 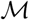 indicates whether solutions were present that include the specified motif (if false, then the motif is disallowed). Column 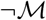 indicates whether solutions were present that do not include the specified motif (if false, then the motif is required). 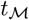 and 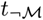 denote the computation times (in seconds) for testing if the motif was disallowed and required. 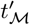 and 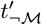 denote the computation times (in seconds) for generating and encoding the motif constraints.

| Motif | $\mathcal{M}$ | $\neg\mathcal{M}$ | $t_{\mathcal{M}}$ ($t'_{\mathcal{M}}$) | $t_{\neg\mathcal{M}}$ ($t'_{\neg\mathcal{M}}$) |
| --- | --- | --- | --- | --- |
| CPFFa | true | true | 13.63(0.09) | 3.32(0.07) |
| CPFFb | true | true | 9.63(0.06) | 9.29(0.07) |
| IFFd | **false** | true | 105.28(0.07) | 8.11(0.17) |
| CNFFa | true | true | 6.89(0.06) | 4.85(0.08) |
| CNFFb | true | true | 1.80(0.06) | 9.79(0.07) |
| IFFa | **false** | true | 122.59(0.08) | 4.72(0.17) |
| IFFb | true | true | 50.49(0.06) | 4.69(0.07) |
| IFFc | **false** | true | 79.28(0.06) | 8.47(0.18) |
| PFB2a | true | true | 4.91(0.07) | 9.30(0.06) |
| PFB2b | true | true | 2.41(0.07) | 7.72(0.06) |
| NFB2 | true | true | 3.06(0.06) | 4.41(0.06) |
| PFB3a | **false** | true | 3.66(0.07) | 1.50(0.07) |
| PFB3b | **false** | true | 9.72(0.06) | 1.63(0.07) |
| NFB3a | **false** | true | 1.62(0.06) | 3.45(0.06) |
| NFB3b | **false** | true | 1.69(0.06) | 2.69(0.06) |

**Table 2.**
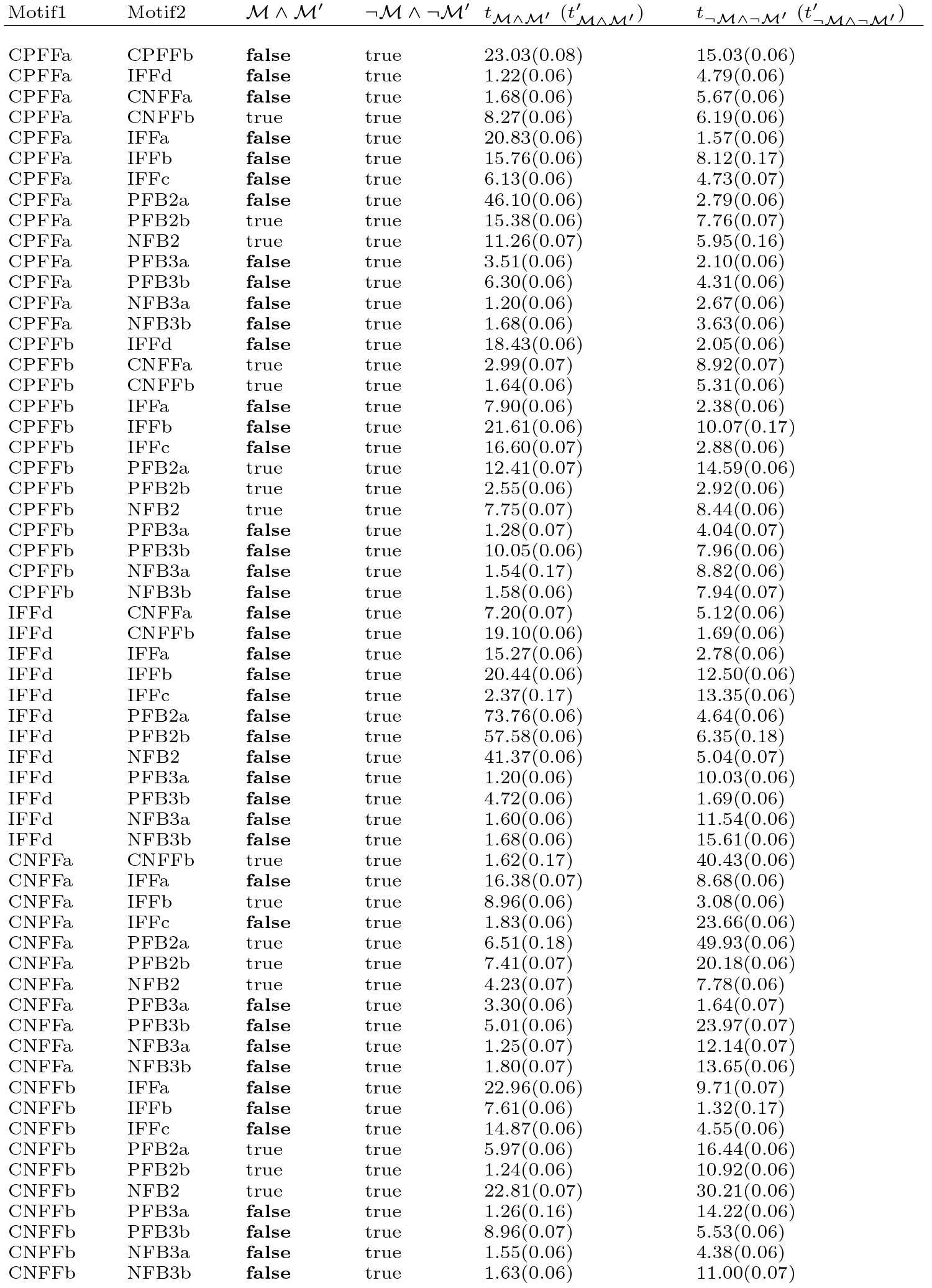
Two-motif analysis results for the 4-step sign-sensitive delay property of the *n* = 3 generic network. Columns are labeled as in Table 1.

**Table 3.**
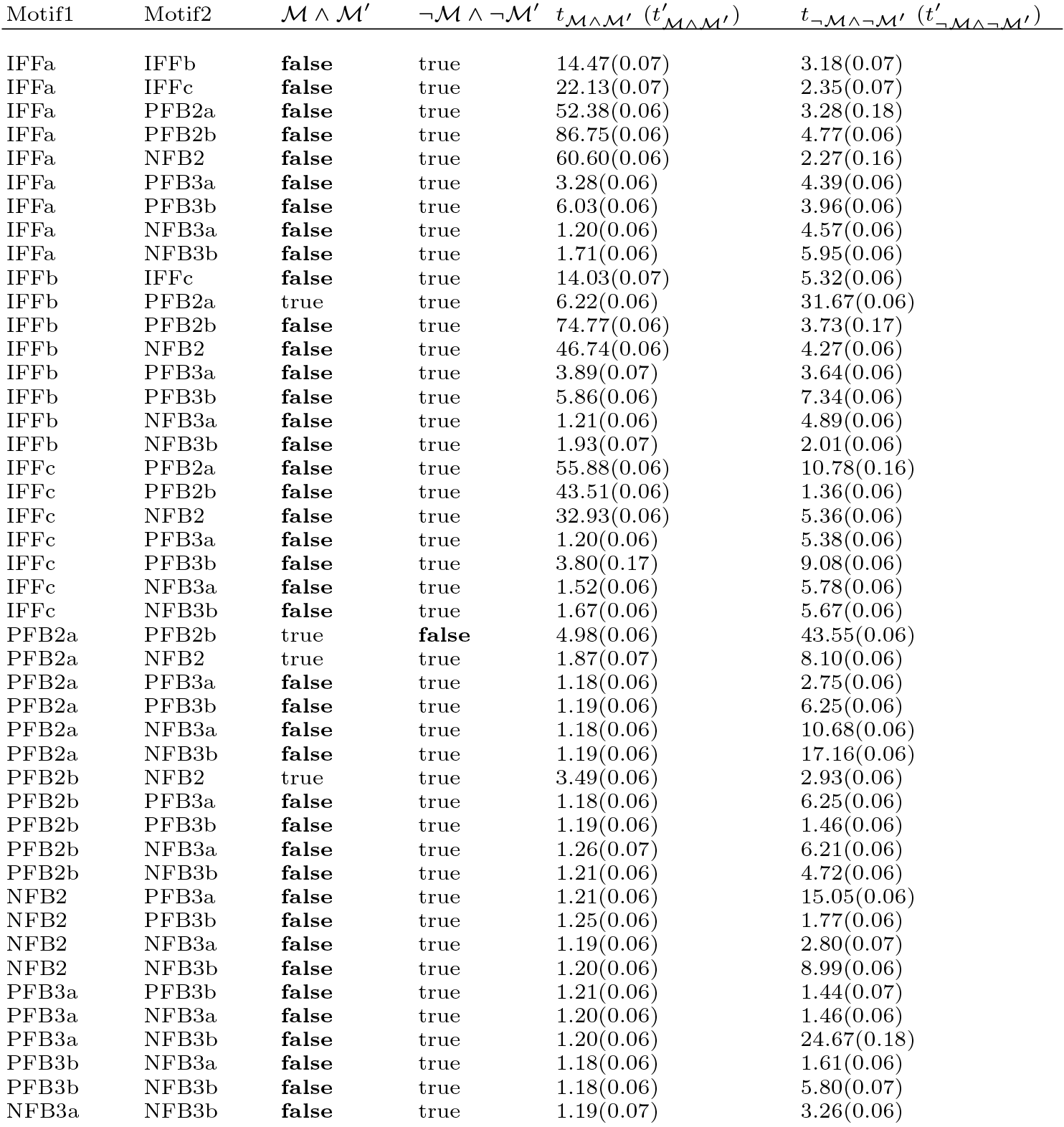
Continued from Table 2.

**Table 4.** Single-motif analysis results for the pulse generation property (pulse width of 4 steps) of the *n* = 3 generic network. Columns are labeled as in Table 1.

| Motif | $\mathcal{M}$ | $\neg\mathcal{M}$ | $t_{\mathcal{M}} (t'_{\mathcal{M}})$ | $t_{\neg\mathcal{M}} (t'_{\neg\mathcal{M}})$ |
| --- | --- | --- | --- | --- |
| CPFFa | **false** | true | 5.86(0.08) | 4.40(0.19) |
| CPFFb | true | true | 3.59(0.07) | 3.75(0.06) |
| IFFd | **false** | true | 8.21(0.06) | 5.50(0.06) |
| CNFFa | **false** | true | 6.09(0.06) | 1.55(0.07) |
| CNFFb | true | true | 3.23(0.06) | 4.17(0.06) |
| IFFa | true | true | 0.92(0.06) | 1.79(0.06) |
| IFFb | true | true | 0.90(0.06) | 5.96(0.07) |
| IFFc | **false** | true | 6.87(0.06) | 1.67(0.06) |
| PFB2a | true | true | 2.50(0.06) | 1.71(0.06) |
| PFB2b | true | **false** | 2.89(0.07) | 11.26(0.06) |
| NFB2 | true | true | 2.14(0.06) | 2.97(0.06) |
| PFB3a | **false** | true | 1.50(0.06) | 3.49(0.07) |
| PFB3b | **false** | true | 2.38(0.06) | 6.26(0.06) |
| NFB3a | **false** | true | 1.36(0.06) | 4.77(0.07) |
| NFB3b | **false** | true | 1.81(0.18) | 1.73(0.06) |

**Table 5.** Single motif analysis results for the stem cell network. Columns are labeled as in Table 1.

| Motif | $\mathcal{M}$ | $\neg\mathcal{M}$ | $t_{\mathcal{M}} (t'_{\mathcal{M}})$ | $t_{\neg\mathcal{M}} (t'_{\neg\mathcal{M}})$ |
| --- | --- | --- | --- | --- |
| CPFFa | true | true | 630.62(19.19) | 460.10(18.65) |
| CPFFb | true | true | 277.47(18.40) | 586.21(18.17) |
| IFFd | **false** | true | 26.27(19.23) | 636.10(19.16) |
| CNFFa | true | true | 489.93(18.43) | 611.28(18.90) |
| CNFFb | true | true | 537.20(18.28) | 493.21(17.90) |
| IFFa | true | **false** | 609.35(19.95) | 540.18(18.39) |
| IFFb | true | true | 568.20(19.15) | 555.86(17.77) |
| IFFc | **false** | true | 515.28(20.74) | 619.15(19.11) |
| PFB2a | true | **false** | 467.51(18.01) | 112.02(18.40) |
| PFB2b | true | true | 505.46(18.21) | 505.40(18.74) |
| NFB2 | **false** | true | 25.05(17.91) | 640.31(17.72) |
| PFB3a | true | **false** | 575.88(17.62) | 525.42(19.04) |
| PFB3b | true | true | 410.93(17.84) | 531.24(17.73) |
| NFB3a | **false** | true | 25.04(17.73) | 572.96(17.65) |
| NFB3b | true | true | 522.14(18.03) | 531.86(17.96) |

## Footnotes

5 While, in general, the motif assignment *θ* is not invertible, *θ*^−1^(*c*) and *θ*^−1^(*c*′) can be defined for the interactions 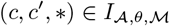 and 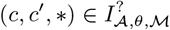

